# A synthesis and meta-analysis on the role of mycorrhizal fungal properties for understanding forest tree functions

**DOI:** 10.1101/2023.03.19.533308

**Authors:** Anis Mahmud Khokon, Ina C. Meier

## Abstract

1. Almost all tree species are associated with mycorrhizal fungi, which help their hosts to acquire soil resources and influence the nutrient economy of the ecosystem. However, despite the fundamental nature of this symbiotic interaction, our understanding of the relationship between fungal properties and host functions is still in its infancy.
2. We compiled a unique database of fungal properties and examine the properties and traits of mycorrhizal fungi in relation to the three main functions of forest trees: resource acquisition, tree production, and carbon release (C).
3. Using quantitative synthesis and meta-analysis, we highlight the current strengths and knowledge gaps regarding tree species, spatial coverage and the relationships between fungal properties and tree functions. Most studies showed that the properties of fungal community assembly, host-symbiont interactions and soil exploration have a positive influence on the resource acquisition and production of host trees. According to a meta-analysis, the composition of the fungal community had a greater influence on resource uptake and tree production than taxon-specific effects. However, some properties of fungal community assembly were negatively related to root carbon (C) exudation (with low certainty), and the influence of these properties and soil exploration on carbon (C) release by host trees through root exudation and litter decomposition remained unclear.
4. We conclude that the influence of mycorrhiza on important tree functions remains an important area for future research, particularly with regard to fungal properties relating to soil exploration and tree properties relating to C release. Such research should progress beyond laboratory measurements of soft traits and increasingly focus on the mycorrhizal fungal properties that are most important for ecosystem functioning.

## 1 INTRODUCTION

Mycorrhizal symbiosis is of fundamental importance to carbon (C) and nutrient cycles in ecosystems, as it facilitates the acquisition of soil resources from host plants in exchange for photosynthates for fungal symbionts. Mycorrhizal roots are found in the majority of all Angiosperm species (Brundrett, 2009) and the symbiosis gained a reputation as a key determinant of terrestrial ecosystem functioning. Approximately 75% of tree species in terrestrial ecosystems associate with arbuscular mycorrhizal fungi (AMF), while the remainder associate with orchid mycorrhizal fungi (OMF), ectomycorrhizal fungi (ECMF) or ericoid mycorrhizal fungi (ERMF; Brundrett & Tedersoo, 2019). Although only 2% of all tree species form the ECM association (van der Heijden et al., 2015), these tree species are widely distributed. As a consequence of multiple evolutionary origins (Pellitier & Zak, 2018) and their wide distribution, ECMF communities are taxonomically more diverse, that is, they are composed of more distantly related fungal species than AMF communities and dominate tree stems in the world’s forests, accounting for around 60% of them (van der Heijden et al., 2015; Steidinger et al., 2019). The nutrient economy of AM and ECM forest ecosystems differs fundamentally (Phillips et al., 2013), reflecting the functional differences between the fungi of these two major mycorrhizal associations. AMF affects the nutrient uptake capacity of tree hosts from competition between root and hyphal uptake only indirectly (Cornelissen et al., 2001). AMF have a high own N demand (Hodge & Fitter, 2010), which AMF can satisfy by scavenging for inorganic nutrients released by saprotrophic microbes (Read & Perez-Moreno, 2003) inter alia from their own labile litter. On the contrary, ECMFs more directly control the uptake of nutrients from their host from the formation of a fungal sheath surrounding fine roots (Tedersoo & Bahram, 2019), which limits nutrient uptake to hyphal tips or non-mycorrhizal root tips, but also protects hosts against soilborne pathogens. ECMF have well-developed saprotrophic capacities and can mine soils for complex N-bearing organic compounds, including their own poor-quality litter (Read & Perez-Moreno, 2003).

The form and properties of root systems are highly plastic and diverse and respond to abiotic and biotic influences. Upon the formation of the symbiosis with mycorrhizal fungi, the trees respond with a change in their fine root properties and functions. When trees form an association with AMF, they increase the cortical area and diameter of the roots to provide an intraradical habitat for their fungal symbionts (Bergmann et al., 2020). In association with ECMF, the morphology and architecture of tree roots change strongly. Trees form root systems with higher branching intensity, that is, they have more root tips per unit length of lower-order roots, providing more root tips for colonization by ECM symbionts and allowing precise foraging in resource-rich soil patches (Liese et al., 2017). In addition, they rely on extensive rhizomorphs (when associated with basidiomycetes) and numerous thin fungal hyphae to explore the soil and exploit the micropores of the soil. Consequently, roots and root-associated fungi can be a major source of long-term C sequestration in ECM forest soils (Clemmensen et al., 2013; Tedersoo & Bahram, 2019). However, despite the multitude of influences of mycorrhizal symbioses on anatomical, morphological, architectural and physiological root traits, and differences in hyphal properties and traits, most investigations of tree-mycorrhizal fungal relationships focus on percentage of root tips or length colonized by mycorrhizal fungi to represent the dependence of trees on the symbiosis (Halbritter et al., 2020; McCormack & Iversen, 2019; Soudzilovskaia et al., 2015).

Plant functional traits are defined as ‘morpho-physio-phenological traits which impact fitness indirectly via their effects on growth, reproduction and survival, the three components of individual performance’ (Violle et al., 2007). Functional traits describe how the morphology and physiology of plants interact with their biotic and abiotic environments and resources (de Bello et al., 2010; Leuschner & Meier, 2018; Violle et al., 2007). They also explain how these characteristics influence the functioning of ecosystems and the provision of ecosystem services. Given the meaning and range of trait-based approaches, functional ecology has received increasing attention in ecological and evolutionary research during the last two decades (Dawson et al., 2021). Compiling trait data into large databases accelerated this development. The *F*ine-*R*oot *E*cology *D*atabase (FRED; Iversen et al., 2017) has recently been developed to compile information on coarse and fine root traits. There is also growing interest in the development of a comparable trait database for fungi, e.g., on the composition of the fungal community composition [FUNGuild (Nguyen et al., 2016); GlobalFungi (Větrovský et al., 2020)], plant biomass responses to mycorrhizal fungal inoculation [MycoDB (Chaudhary et al., 2016)], fungal ecology [FUN^FUN^ (Zanne et al., 2019)], functional properties of fungi [FungalTraits (Põlme et al., 2020)], or on mycorrhizal associations [FungalRoot (Soudzilovskaia et al., 2020)].

Mycorrhizal properties and traits can be influenced by both the plant host and the associated community of fungal symbionts, which can have considerable variation in morphological, physiological, and biochemical traits (Chaudhary et al., 2022; Hazard & Johnson, 2018). This bilateral influence on mycorrhizal properties and traits complicates their analysis. More complex fungal communities are profitable to the host, as they increase the chance of greater functional diversity among the associated mycorrhizal fungi (Baxter & Dighton, 2001; Khokon et al., 2023; Köhler et al., 2018). For example, ECMF communities composed of fungal symbionts with various types of exploration (distance of soil exploration from the host tree as based on morphological traits of colonized root tips and emanating hyphae; Agerer, 2001) have higher chance of spatial and functional complementarity among the emanating fungal mycelia. For decades, fungal ecologists have used selected mycorrhizal fungal traits, such as mycorrhizal colonization, to understand host and ecosystem functioning (Agerer, 2001; Chaudhary et al., 2022; Fernandez & Kennedy, 2018; Lekberg & Helgason, 2018; Zanne et al., 2019). However, a holistic examination of mycorrhizal fungal properties or traits that are decisive for major tree functions or mycorrhizal economics remains elusive (*cf.* Bergmann et al., 2020; Wright et al., 2004). Here, we compile global quantitative data on the properties and characteristics of mycorrhizal fungi at both at the individual and community levels by conducting an extensive literature review. We then relate these properties to the major functions of forest trees, categorized as ‘resource acquisition’, ‘tree production’ and ‘C release’ (via root exudation and litter decomposition). Drawing on our expert knowledge, we summarized the underlying rationales and conceptualized potential relationships between fungal properties and major functions of forest trees (see Leuschner & Meier, 2018). Finally, we tested these conceptual models in a meta-analysis using the assembled database. Here, we critically evaluate current strengths and gaps in our knowledge of the relationships between the properties of fungal symbionts and host trees, and highlight avenues for future research that could improve our understanding of how trees respond to limited soil resources and affect soil C cycling in forest ecosystems.

## 2 MATERIALS AND METHODS

### 2.1 The mycorrhizal fungal trait database

We assembled information on mycorrhizal fungal properties and traits used to understand forest tree functions from a systematic literature survey using the Web of Science (Clarivate Analytics, Philadelphia, PA, USA). We focus on six forest tree functions in the categories (i) resource acquisition (N, P and water uptake), (ii) plant productivity (merged above-ground and below-ground data), and (iii) plant C release (root exudation and litter decomposition), for which we used the following search term: “(mycorrhiza* OR ectomycorrhiza*) AND (nitrogen OR phosphor* OR water OR produc* OR decompos* OR exud*) AND (tree*) NOT (grass*) NOT (herb*)”. The survey included references between 1986 and September 20, 2022 (date of the first study found and date at which the survey was carried out, respectively) and covered nearly 2,900 published studies on the mycorrhizal fungal properties and traits of forest trees (Figure S1.1). Subsequently, we screened the initial literature list in accordance with the guidelines ‘Preferred Reporting Items for Systematic Review and Meta-Analyzes’ (PRISMA) [https://www.prisma-statement.org; (Moher et al., 2010; Page et al., 2021)]. All non-English, duplicates, and review articles (to avoid hidden duplicates) were excluded from the records. We checked all titles and abstracts for the eligibility criterion, that is, relevant information on mycorrhizal fungal properties and traits in relation to tree functioning. For the remaining articles, we also assessed whether the full text met the eligibility criteria (e.g., only original and peer-reviewed studies; woody plants as target species; and the absence of confounding treatments; Figure S1.1). As a result, a total of 464 studies entered the subsequent quantitative synthesis and meta-analysis. For each selected publication, we collected information on the investigated tree species, tree function, tree organ, type of association (AM, ECM, or ERM) and mycorrhizal fungal properties/traits, study type (controlled experiment or field observation), the significance and type of the functional response of the tree for a given mycorrhizal fungal property/trait (positive, negative, or neutral), location of the study and additional general information from the imprint of the printer.

For the meta-analysis, we collected the following data on our variable of interest when reported: means, correlation coefficient (*r*), *F statistics, t statistics,* sample size (*n*), standard deviation (*SD*), standard error (*SE*) and 95% confidence interval (*CI*). Unspecific error bars were recorded as SE. Studies were excluded if they provided only mean values without associated descriptive statistics. Studies involving mycorrhizal associations often lack a defined control group and, therefore, AM associations were treated as control, and ECM associations as treatment. During data extraction, we also applied consistent decision rules to ensure comparability between experiments. Where multiple treatment levels were reported, we selected the treatment most representative of typical field conditions or least likely to introduce confounding stress, unless all levels were required for independent analysis. Figures were digitalized using WebPlotDigitizer 4.1 software (https://automeris.io/). Finally, our data set for meta-analysis consisted of 3,451 observations from 353 publications for mycorrhizal properties/traits at the community level, and 2488 observations from 177 publications for the mycorrhizal fungal taxon-specific data set.

### 2.2 Replication statement

Scale of inference: Tree species

Scale at which the factor of interest is applied: Tree species

Number of replicates at the appropriate scale: 1-256 observations (Table 2) from 464 selected research articles (*cf.* PRISMA protocol; Figure S1.1)

**TABLE 1.** Frequency of studies investigating properties/traits of mycorrhizal fungi (at the individual or community level) in relation to major functions of forest trees (i.e., resource acquisition, tree production, or carbon release) under controlled experimental or natural field conditions between 1986 and 2022.

|  | Controlled<br>experiment | Field<br>observation | Total |
| --- | --- | --- | --- |
| ***Fungal community assembly*** |  |  |  |
| Genetic identity | **225** | 44 | 269 |
| Community composition | **73** | 28 | 101 |
| Genetic richness | **32** | 9 | 41 |
| Genetic diversity | 3 | 3 | 6 |
| Genetic evenness | 0 | 1 | 1 |
| ECM functional genes | 0 | 1 | 1 |
| ***Host-symbiont interaction*** |  |  |  |
| Mycorrhizal association type | 18 | **52** | 70 |
| Colonization intensity | 24 | 23 | 47 |
| ECM tree dominance | 0 | **11** | 11 |
| Common ECMF network | 0 | **4** | 4 |
| Hyphal proportion per plant | 1 | 0 | 1 |
| ***Soil exploration*** |  |  |  |
| AMF spore density | **3** | 2 | 5 |
| AMF hyphal length | 1 | 0 | 1 |
| ECMF exploration type | 0 | **3** | 3 |
| ECM mycelial diameter | 1 | 0 | 1 |
| Hyphal foraging precision | 1 | 0 | 1 |
| ECM mycelium density | 1 | 0 | 1 |

**TABLE 2.**
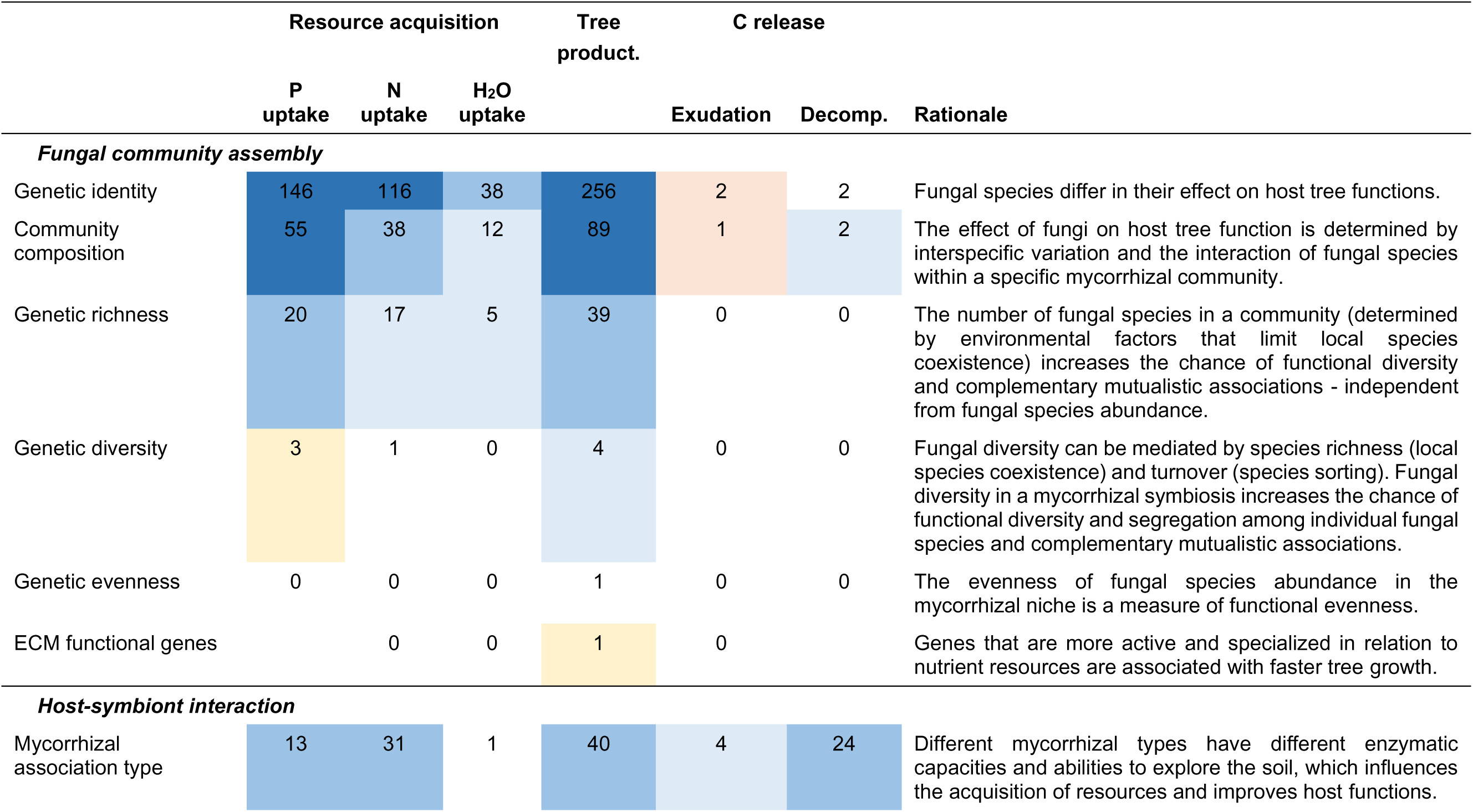
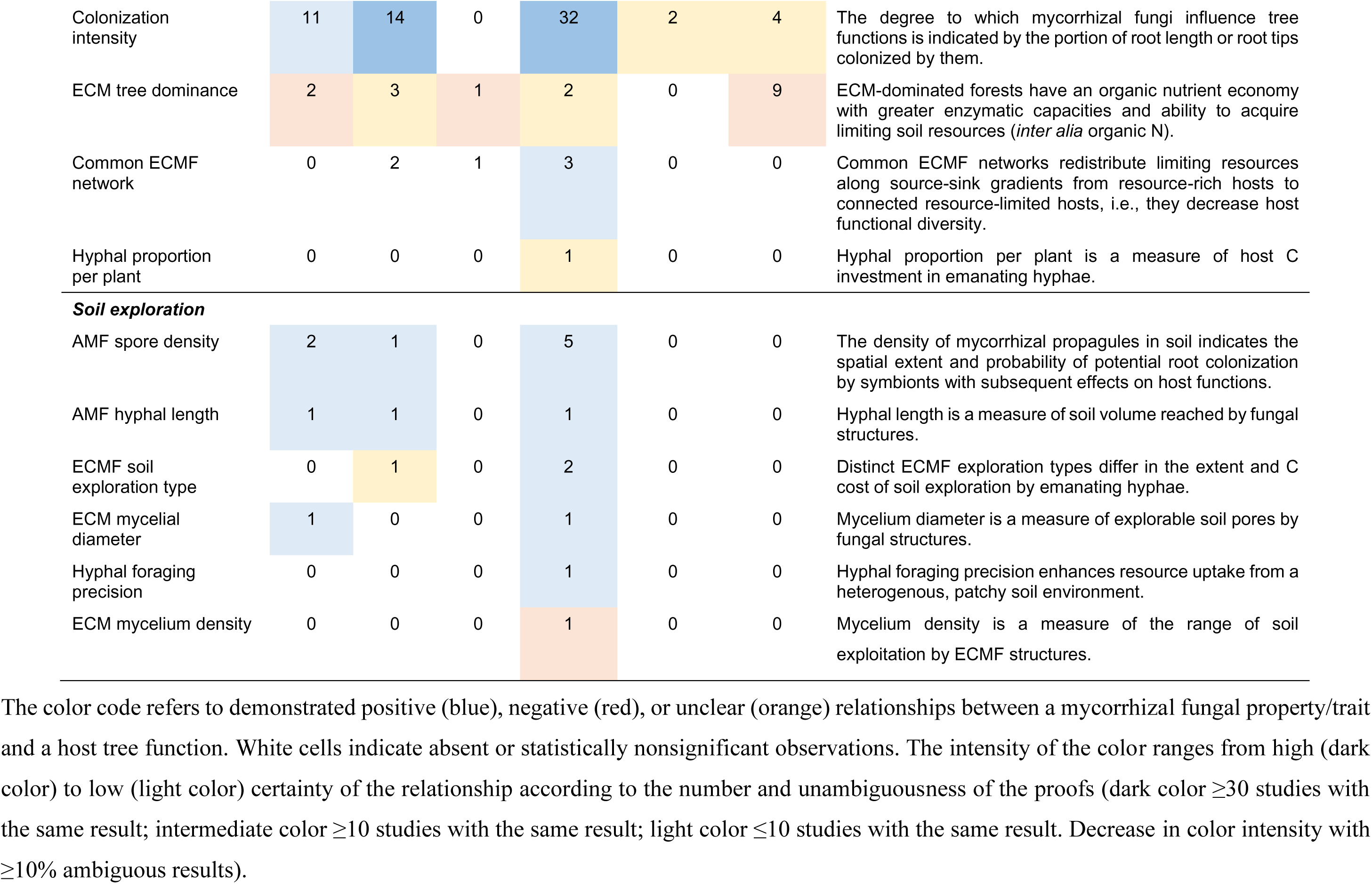
Frequency of observed relationships between the properties/traits of mycorrhizal fungi (at the individual or community level) and major functions of forest trees (i.e., resource acquisition, tree production and carbon release) between 1986 and 2022.

**TABLE 3.**
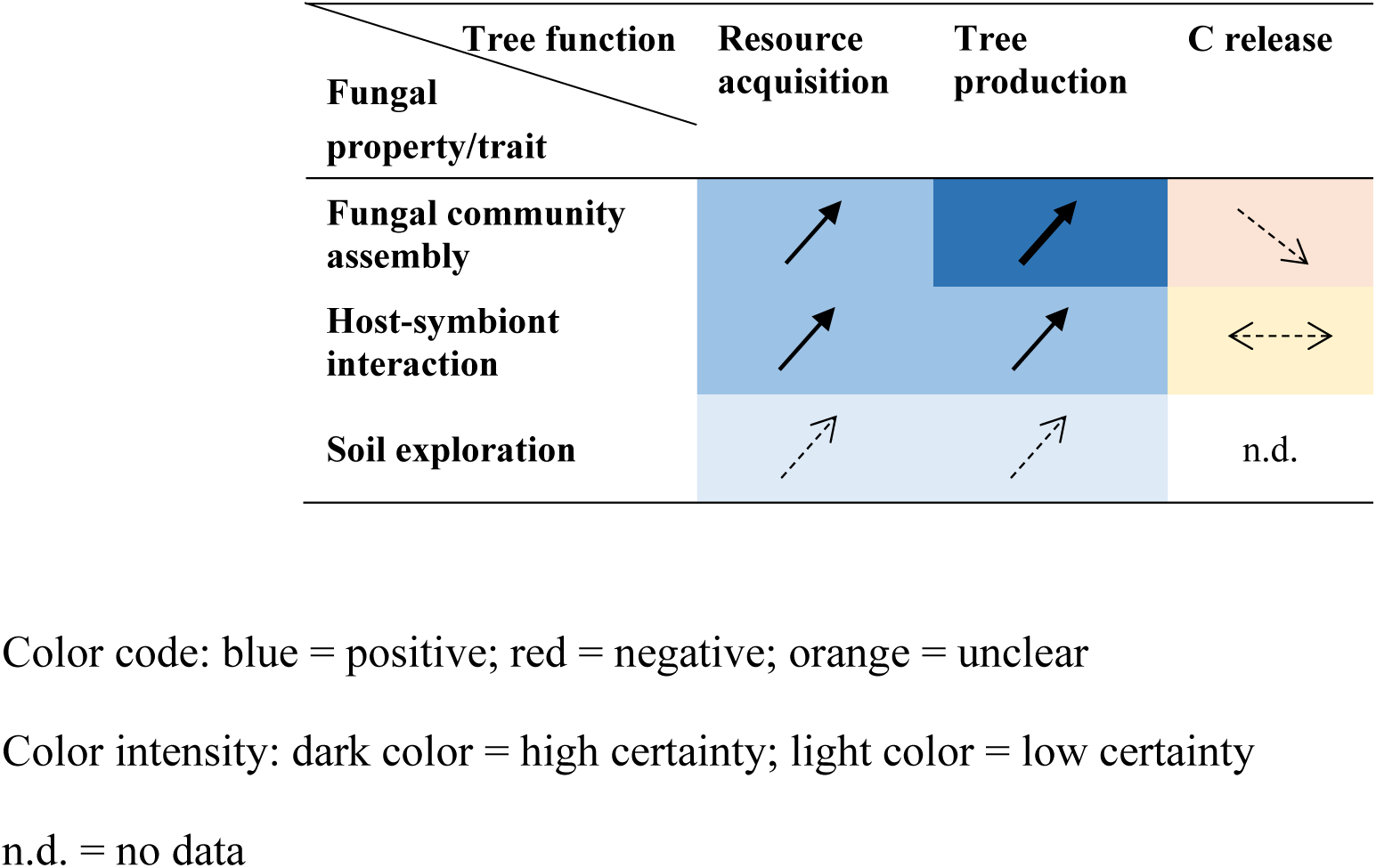
Summary of the observed relationships between the properties/traits of mycorrhizal fungi (at the individual or community level) and major functions of forest trees (i.e., resource acquisition, tree production and carbon release) between 1986 and 2022 (based on a systematic literature review; *n* = 464 studies).

### 2.3 Data preprocessing

When the study reported *SE*, the corresponding *SD* was calculated by multiplying *SE* by the square root of the sample size *n* (see Tang et al., 2023). When the study reported the *CI*s, the corresponding *SD* was calculated by multiplying the square root of *n* by the difference between the upper and lower *CIs* and dividing the result by two times 𝑍_𝛼/2_, where 𝑍_𝛼/2_represents the Z-value for the corresponding significance level (e.g., 1.96 at *α* = 0.05). When mean values were unavailable (for example, when data came from box plots), the sum of the first quartile (*Q1*), the median (*Q2*), and the third quartile (*Q3*) was divided by three. The interquartile range (*IQR*) was calculated from the difference between *Q3* and *Q1*. The *SD* was then estimated from the *IQR* using the following small-sample, bias-corrected formula (Wan et al., 2014):

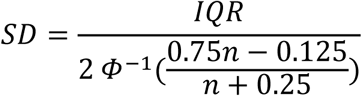

Here, *Φ* denotes the cumulative distribution function (CDF) of the standard normal distribution.

### 2.4 Effect size and variance calculation

To determine the effect sizes for the influence of mycorrhizal fungal properties/traits on tree functions, the standardized mean differences (Hedges’ *g*) and their variances were calculated using the *escalc()* function (*measure = ‘SMD’) of* the *metafor* package in *R* (*v4.8.0*; Viechtbauer, 2010) in R, when means, sample sizes (𝑛), standard deviations (*SD*), or *t*-statistics were available for both control and treatment groups. As *metafor* does not directly compute Hedges’ *g* and variance from correlation coefficients (𝑟) or *F*-statistics, we applied the following conversions. When *r* was reported, Cohen’s *d* was calculated as (Borenstein et al., 2009; Rosenthal, 1991):

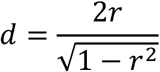

Cohen’s *d* was adjusted to Hedges’ *g* using small-sample correction factors (*j*):

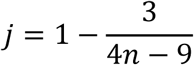

The effect size (Hedges’ *g*) was calculated as follows:

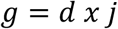

The variance of *r* (*Vr*), Cohen’s *d* (*Vd*) and Hedges’ *g* (*Vg*) were calculated as follows:

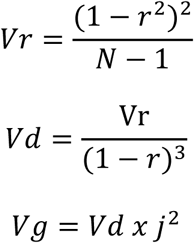

When *F*-statistics, sample size (*n*) for control (*nc*) and treatment (*nt*) groups were reported, Cohen’s *d* was calculated as (Borenstein et al., 2009; Hedges & Olkin, 1985):

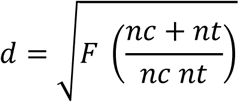

Cohen’s *d* was adjusted to Hedges’ *g* using the correction factors (*j*) (Hedges & Olkin, 1985):

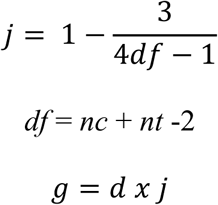

The variance of Cohen’s *d* (*Vd*) and Hedges’ *g* (*Vg*) was calculated as:

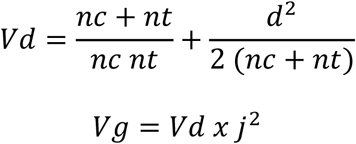

The effect size (Hedges’ *g*) and the variance were calculated for each observation associated with mycorrhizal fungal properties/traits and tree functions. The impact of mycorrhizal properties/traits on tree functions was evaluated based on the Hedges’ *g* statistic. A positive effect was indicated when g > 0, and a negative effect was indicated when *g* < 0 (Marro et al., 2022). The mean effect sizes were deemed statistically significant when the overall estimate (95% *CI*) range did not overlap zero (Borenstein et al., 2009).

To reduce statistical dependence and avoid overrepresentation of studies reporting multiple results (i.e., effect sizes) for the same combination of property/trait and function, effect sizes and variance were aggregated at the level of ‘*study x mycorrhizal trait (and taxon-specific) x tree function’*. For each unique combination, individual effect sizes were pooled using a fixed effect model with inverse-variance weighting, implemented via the *rma.uni()* function in the *metafor* package in *R* (Viechtbauer, 2010). The robustness of the aggregation approach was assessed by comparing fixed-effects and random-effects pooling within a random subset of the *study × mycorrhizal trait (and taxon-specific) × tree function*. The differences in the pooled estimates were minimal (mean absolute difference = 0.27 and median ∼0 for the mycorrhizal traits dataset; mean absolute difference = 0.69 and median ∼0 for the taxon-specific mycorrhizal fungal data set), thus supporting the use of fixed effects models for the aggregation within the study.

During the data cleaning, we removed seven extreme effect sizes *(|yi|* >10), which were mainly attributed to combination of large mean values, very small *SD,* and small sample size. Furthermore, effect sizes based on fewer than two observations and/or derived from a single article were further excluded from analyses to avoid effects by the small sample size (Borenstein et al., 2009). At this stage, eighteen observations from sixteen publications were removed. The same filtering criteria were applied when analyzing the influence of taxon-specific effects of AM and ECM on tree functioning. The final data set comprised 714 observations for mycorrhizal fungal properties/traits and 487 observations for taxon-specific effects on tree functioning.

### 2.5 Visualization of the spatial distribution of data

Data were visualized with the *ggplot2* package (Wickham, 2009) in *R*, version 4.2.2 (R development Core Team., 2020). The publicly available global geographic data set with information on longitudes, latitudes, and regions was imported into *R* using the map_data(“world”) function prior to creating the global map. The Antarctica region was excluded from the global map by implementing the *filter (region! = ‘Antarctica’* function utilizing the *tidyverse* package (Wickham et al., 2019), given the absence of any published study in the Antarctic region. Furthermore, our mycorrhizal trait dataset was merged with the global geographical dataset and sorted according to the city name using the *left_join (by=“region”)* function to create the global map. The different color gradients were used from the *viridis* package (Garnier et al., 2021) to demonstrate the aggregate number of studies on the global map.

### 2.6 Phylogenetic relatedness of the investigated tree species

The construction of a phylogenetic tree was achieved by generating its backbone using the *phyl.maker()* function from the *V.Phylomaker* package in *R* (Jin & Qian, 2019), and subsequently visualizing the tree using the *ggtree()* function from the *ggtree* package (Yu et al., 2017). We annotated the phylogenetic tree using the *geom_text2()* function to label the node numbers, the *geom_tiplab2()* function to align the tree species, and the *geom_hilight() and geom_cladelabel()* function to highlight the most dominant genera from the *ggtree* package (Yu et al., 2017).

### 2.7 Publication bias

Publication bias was assessed on the results using several approaches in the *metafor* package in *R* (Viechtbauer, 2010). Initially, funnel plots were subjected to visual inspection by plotting Hedges’ *g* against *SE*, followed by further evaluation of funnel plot asymmetry through Egger’s regression test using the *regtest()* function (Egger et al., 1997; Tang et al., 2023). In instances where such disparities were identified, the prospective influence of unpublished studies on the conclusions derived was subsequently evaluated by calculating Rosenberg’s fail-safe number utilizing the *fsn()* function (Rosenberg, 2005; Marro et al., 2022). A failsafe number is considered robust when a value of ‘fsn’ is five times higher than the total number of publications plus ten (Rosenthal, 1991; Marro et al., 2022). Finally, the sensitivity of the results was explored graphically by performing a leave-one-out analysis using the *leave1out()* function (Tang et al., 2023).

### 2.8 Data analysis

For quantitative synthesis, in the first step, we identified significant positive or negative responses of mycorrhizal properties/traits to tree functions as well as absent or nonsignificant relationships as published in the original publication. In the next step, we calculated the relative proportion of a specific response of a mycorrhizal property/trait to a tree function across all tree organs of each study separately. The response was defined as unambiguous positive or negative, when the relative proportion of the response was ≥60% of all responses, and as unclear, when the relative proportion of the response was ≤60%. We defined the certainty of the relationship between a mycorrhizal property/trait and a tree function as high, when ≥30 studies had the same result, and as intermediate and low certainty, when ≥10 and ≤10, respectively, had the same result. We decrease the level of certainty when ≥10% of the studies were ambiguous.

For meta-analysis purposes, the mean effect sizes were visually represented using forest plots with the *metafor* package in *R* (Viechtbauer, 2010). Quantification of community-specific and taxon-specific fungal effects on each tree function across studies was achieved by employing mixed-effects meta-regression models (Marro et al., 2022). In the models considered, heterogeneity among effect sizes was calculated using *Cochran’s Q* statistic (Borenstein et al., 2009; Marro et al., 2022). In this approach, *QE* and *QM* represent residual heterogeneity and heterogeneity explained by moderators (i.e., mycorrhizal property/traits or taxa). All models were fitted with the *rma()* function in R (Viechtbauer, 2010).

## 3 RESULTS

### 3.1 Global and tree species representation of the relationship between mycorrhizal fungal properties and host tree functions

Most studies relating mycorrhizal fungal properties to forest tree functions between the study years 1986 and 2022 were conducted on the European continent (125 studies) and in two countries of the Northern Hemisphere, which are dominated by the temperate climate zone (USA: 65 studies, China: 63 studies; Figure 1). These focal areas for mycorrhizal trait studies are followed by three countries dominated by tropical climate (that is, Brazil, India, and Senegal), in which each 19 to 26 studies were conducted in the same time span. Only a few studies of mycorrhizal trait studies were published internationally for three countries belonging to the largest countries of the world (Canada: 13 studies; Australia: 10 studies; Russia: 1 study). The gray spots on the global map of studies relating mycorrhizal fungal propertys to forest tree functions remain in large parts of hot-dry or tropical Africa, continental Central Asia, and in (sub) polar Greenland (Figure 1).

**FIGURE 1.**
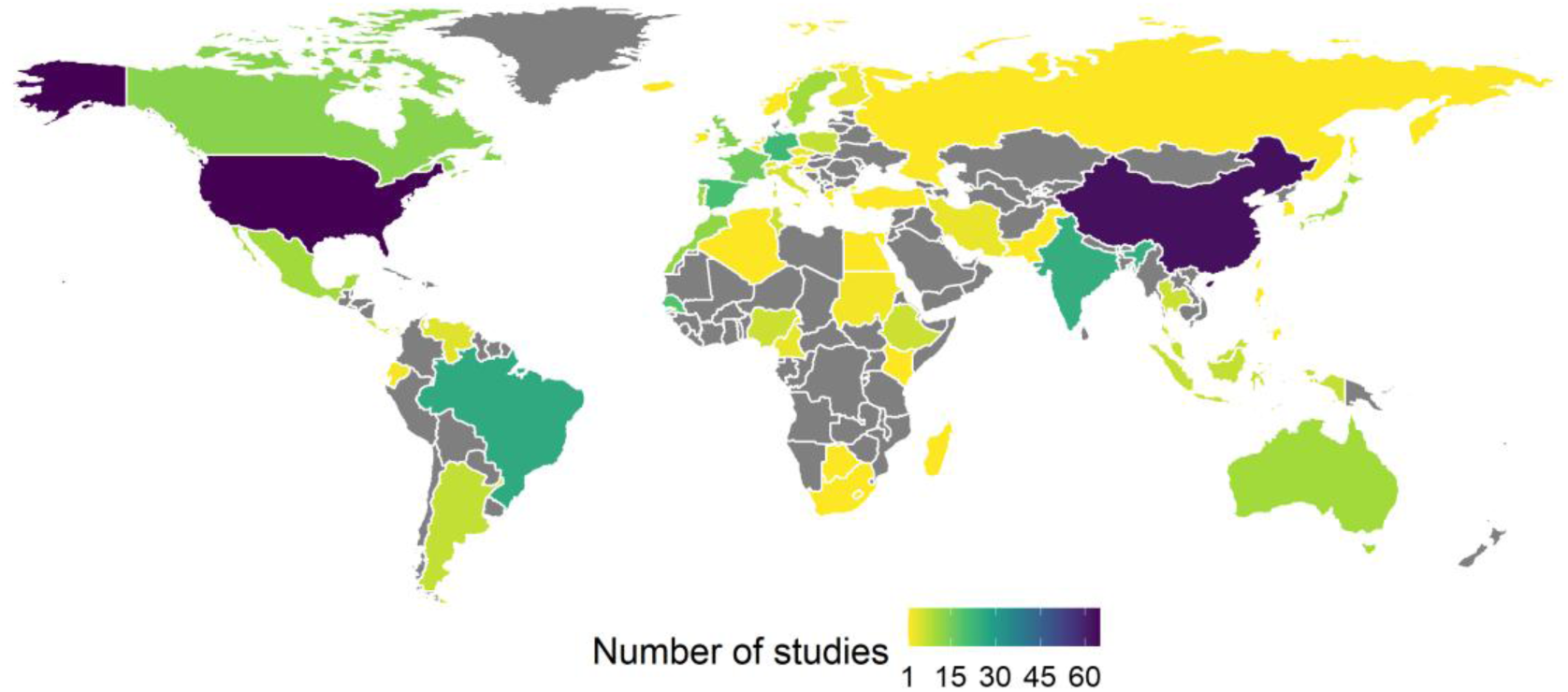
Global distribution of studies investigating properties/traits of mycorrhizal fungi in relation to major functions of forest trees between 1986 and 2022 (*n* = 464 studies).

A total of 501 tree species were used in 464 publications to relate mycorrhizal fungal properties to forest tree functions between the study years 1986 and 2022 (Figure 2). The most abundant tree species used in these publications was the economically important fruit tree olive (*Olea europaea* L.), which is followed in number by sugar maple (*Acer saccharum* Marshall), European beech (*Fagus sylvatica* L.), tulip tree (*Liriodendron tulipifera* L.) and Scots pine (*Pinus sylvestris* L.) (Table S1.1). More than 200 studies investigated trees of the Fabaceae family, which were represented by almost 100 tree species. Fewer studies were conducted in the pine and beech families, which were observed by fewer tree species (*c.* 150 and 110 studies, respectively; Pinaceae: 44 tree species; Fagaceae: 42 tree species). However, in a comparison of the studied genera, *Pinus* (Pinaceae) and *Quercus* (Fagaceae) were the most widely used genera, followed by *Acer* and (with distance) *Betula* and *Fagus*. Among these dominantly investigated genera, the genus *Pinus* represents ECM trees with evergreen leaf habit (Figure 2). Pine tree species are phylogenetically the most distant from the other genera that were investigated in a dominant way, since they were the only gymnosperms among these. Different successional statuses of deciduous ECM trees were represented by *Betula* (early successional), *Quercus* (mid successional), and *Fagus* (late successional), which are all Fagaceae and thus phylogenetically closely related. The genus *Acer* (Sapindaceae), which is located in the middle of the phylogenetic tree, was often used as representative of AM tree species of mid/late successional status. The less frequently studied genera *Olea* and *Acacia* represent more special cases of evergreen AM tree species (that is, *Olea europaea*) and evergreen AM/rhizobia tree species (*Acacia*), respectively. *Olea* is phylogenetically closely related to the genus *Fraxinus* (Oleaceae), which was used as a representative of early / mid successional tree species of AM.

**FIGURE 2.**
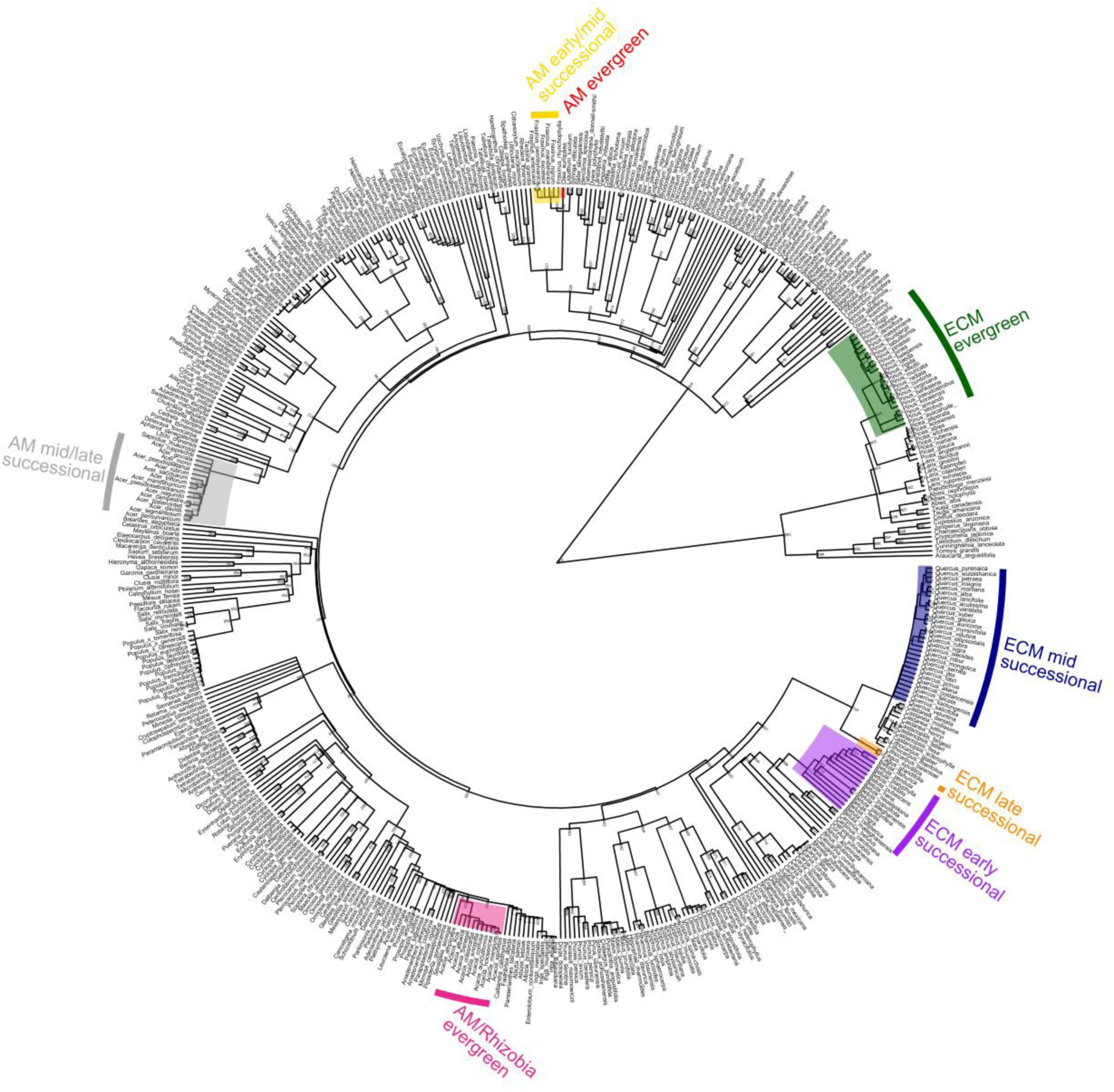
Phylogenetic relationship between the tree species studied to investigate the influence of the properties/traits of mycorrhizal fungi on major functions of forest trees between 1986 and 2022. The nodes representing the genera that dominated the investigations are highlighted in different colours. Color code (usage of tree species in order): green = *Pinus*, blue = *Quercus*, grey = *Acer*, purple = *Betula*, orange = *Fagus*, pink = *Acacia*, yellow = *Fraxinus*, red = *Olea*. Mycorrhizal types: AM = arbuscular mycorrhiza, ECM = ectomycorrhiza.

Most studies investigating ECM mycorrhizal properties/traits were carried out in Europe and the Global North (39% and 89%, respectively), which also accounted for the largest number of studies (423 out of 550; Figure 3). Studies on AM properties were more evenly distributed, with a significant proportion taking place in Asia (33% of all studies). The majority of studies on mycorrhizal properties/traits in Africa and South America focused on AM symbiosis. Only a few studies, mainly in Europe, investigated ERM properties/traits (six studies comparing ERM and ECM). Almost all studies comparing the properties and traits of the two major types of mycorrhizal association originate from the Northern Hemisphere, with many from North America (33 studies).

**FIGURE 3.**
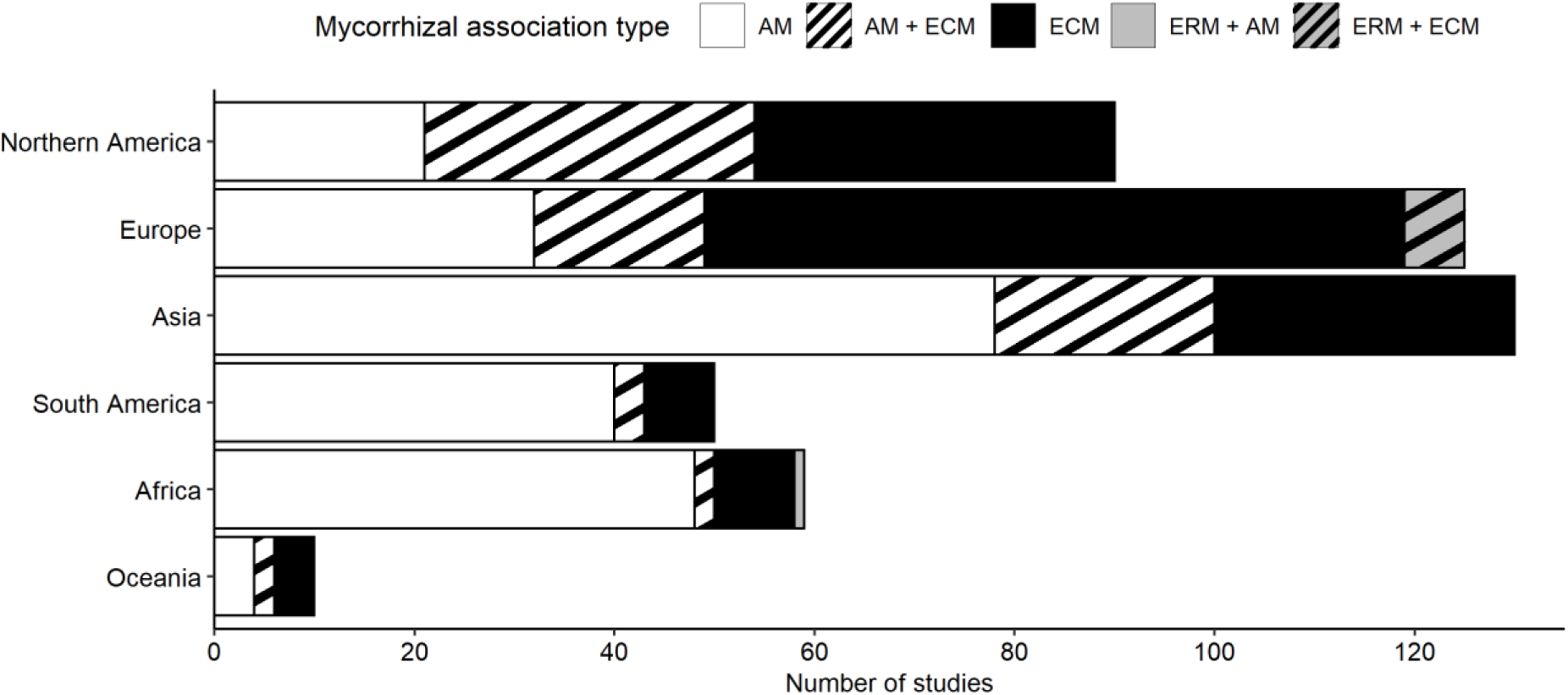
Continental distribution of the different types of mycorrhizal association that were studied in order to investigate the properties/traits of mycorrhizal fungi in relation to major functions of forest trees between 1986 and 2022. Mycorrhizal types: AM = arbuscular mycorrhiza, ECM = ectomycorrhiza, ERM = ericoid mycorrhiza.

### 3.2 Mycorrhizal fungal properties used to understand the functioning of forest trees

Our literature survey resulted in 17 mycorrhizal fungal properties in the categories (i) fungal community assembly, (ii) host-symbiont interaction, and (iii) soil exploration, which were analyzed to understand six target forest tree functions (Table 1). Most of the studies of mycorrhizal functions focused on the influence of the fungal community assembly on tree functions (419 studies), primarily by analyzing the effect of the genetic identity of mycorrhizal fungi, the composition of the fungal community, or the richness of the mycorrhizal fungal species. Almost 80% of all studies on the influence of the fungal community assembly on tree functions were conducted under controlled experimental conditions, and only one fifth in natural field settings.

Fewer studies (*n* = 133; Table 1) examined properties relating to host-symbiont interaction. Of these, over 60% focused on mycorrhizal association type and ECM dominance as fungal properties, primarily through field observations. Only one third of the studies on host-symbiont interactions focused on colonization intensity in experimental or field studies. The USA was the global leader in field observations of host-symbiont interactions (Figure S1.2).

Fewer than 2% of all mycorrhizal studies have focused on soil exploration by mycorrhizal hyphae (see Table 1). Six experimental studies considered arbuscular mycorrhizal fungus (AMF) spore density and hyphal length as factors influencing tree function, and three field observations tested the effect of a specific ectomycorrhizal fungus (ECMF) exploration type. Overall, little attention was paid to the potentially significant impact of fungal diversity, functional genes and common mycorrhizal networks on host tree functions.

### 3.3 Mycorrhizal fungal properties and taxon-specific effects on forest tree functioning

In general, mycorrhizal fungal properties had significant positive effects on N uptake (*g* = 0.74, 95% *CI* [0.14, 1.34], *QM* = 20.20, *p* = 0.001; Figure 4a), P uptake (*g* = 0.98, 95% *CI* [0.36, 1.60], *QM* = 16.50, *p* = 0.01; Figure 4b) and plant productivity (*g* = 0.89, 95% *CI* [0.37, 1.41], *QM* = 27.83, *p* = 0.001; Figure 4c). These positive responses were primarily driven by the composition of the mycorrhizal fungal community, the colonization intensity, the genetic richness and the genetic identity (Figure 4a–c). On the contrary, the mycorrhizal association type and ECM tree dominance showed negligible effects on N uptake and productivity, and the decomposition tended to decline under ECM tree dominance (*g* = –1.36, 95% *CI* [–2.00, – 0.72]; Figure 4d).

**FIGURE 4.**
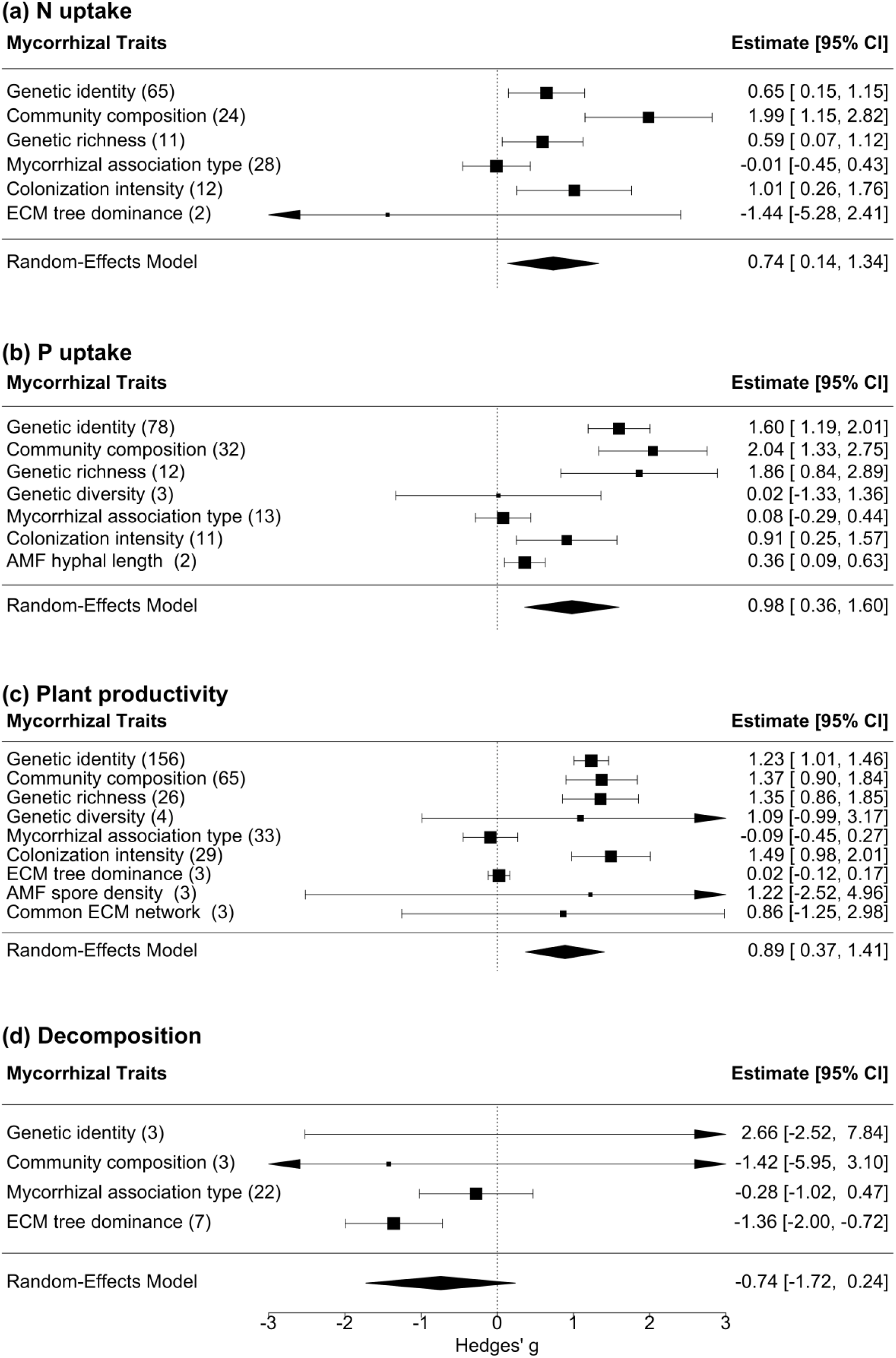
Mean effect sizes (Hedges’ *g*) and estimates of the random effects model (95% confidence intervals) calculated for the influence of the properties/traits of mycorrhizal fungi on major functions of forest trees: **(a)** plant nitrogen (N) uptake, **(b)** plant phosphorus (P) uptake, **(c)** plant productivity and **(d)** decomposition. The frequency of observations (*n*) is given in brackets behind the mycorrhizal fungal property/traits and is represented by the size of the squares showing the mean effect sizes. Significant effect sizes do not overlap with the null effect (*g* = 0).

At the taxon-specific level, both AM and ECM fungal species showed generally positive but statistically nonsignificant effects on N uptake (*g* = 0.70, 95% *CI* [0.18, 1.22], *QM* = 22.18, *p* = 0.33; Figure S1.3), P uptake (*g* = 1.51, 95% *CI* [1.16, 1.86], *QM* = 32.59, *p* = 0.30; Figure S1.4), and plant productivity (*g* = 1.13, 95% *CI* [0.90, 1.37], *QM* = 49.08, *p* = 0.39, Figure S1.5). Several AM taxa, including *Rhizophagus intraradices*, *R. irregularis*, *Funneliformis mosseae*, and *Glomus aggregatum*, as well as ECM taxa such as *Cenococcum geophilum*, *Hebeloma mesophaeum*, *Helvella lacunosa*, *Suillus grevillei*, and *Laccaria laccata*, exhibited consistently positive associations with nutrient uptake and productivity (Figure S1.3-5). These results indicate that there are no systematic differences among AM and ECM taxa, but the overall pattern reveals broadly positive and variable taxon-specific responses, suggesting that both AM and ECM contribute to enhanced nutrient uptake and plant productivity through distinct but functionally convergent mechanisms.

### 3.4 Publication bias and sensitivity analysis

According to an Egger’s test, there was no significant publication bias in the effect of mycorrhizal fungal properties on tree N uptake (p > 0.05; Table S1.2, Figure S1.6). However, asymmetries were evident in funnel plots of other tree functions (P uptake, plant productivity, decomposition; Figures S1.6, S1.8). However, the Rosenberg’s failsafe numbers were greater than the robustness threshold of 5k + 10 (Table S1.2; see Tang et al., 2023), except for water uptake and exudation. The latter were therefore excluded from the meta-analysis. Furthermore, exudation and decomposition variables were excluded from the taxon-specific dataset for meta-analysis due to insufficient data (Table S1.2). According to sensitivity analyzes, no singular publication controlled our results (Figure S1.7; Figure S1.9).

### 3.5 Concept of relationships between fungal properties and tree functioning

Drawing on existing knowledge, we have developed a conceptual framework that delineates the relationships between the effects of mycorrhizal fungal properties and the key functions of host trees (resource acquisition, tree production, and C release). This framework was separated for properties related to the category ‘fungal community assembly’ (Figure 5a) and properties related to the categories ‘host-symbiont interaction’ and ‘soil exploration’ (Figure 5b), respectively. The conceptual framework is based on three widely assumed effect-response relationships: (i) More complex fungal communities with greater fungal species richness and/or species evenness and greater species diversity enhance host tree functioning with respect to soil resource acquisition and C release from root exudation and decomposition (Baxter & Dighton, 2001; Clemmensen et al., 2021; Khokon et al., 2023; Köhler et al., 2018). (ii) ECMFs with different exploration types differ by their mycelium density and hyphal foraging precision (Agerer, 2001). Greater soil exploration by mycorrhizal hyphae (i.e. hyphal foraging precision at reduced mycelium density) increases resource acquisition and production of plants (Lilleskov et al., 2011, 2019). (iii) Mycorrhizal colonization of tree roots exerts a control on root C release through exudation or decomposition (Argiroff et al., 2022; Fanin et al., 2022; Meier et al., 2013; Tedersoo & Bahram, 2019). We tested this conceptual framework against the published results on relationships between mycorrhizal fungal properties and key functions of host trees.

**FIGURE 5.**
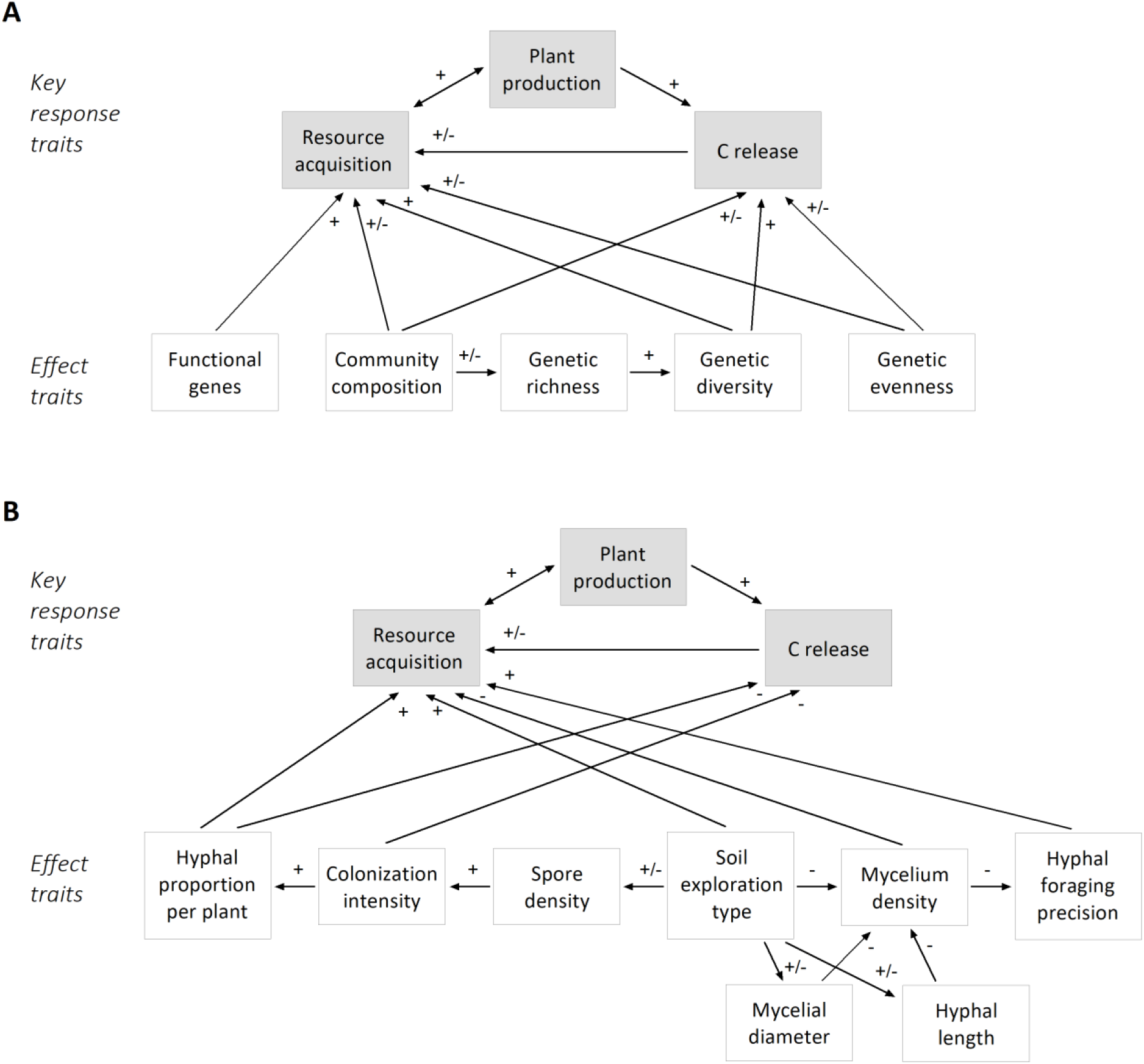
Conceptual models of the plausible relationships between the properties/traits of mycorrhizal fungi (**a**: fungal community assembly; **b**: host-symbiont interactions and soil exploration) and the major functions of forest trees. ’+’ refers to a positive effect and ‘-’ refers to a negative effect.

The published results for the relationships between mycorrhizal fungal properties and tree function demonstrate an unambiguous positive influence of fungi of a specific genetic identity, a specific community composition and genetic richness on properties related to soil resource acquisition and tree production (Table 2, 3; Figure 4). The few existing studies on fungal genetic diversity found a positive influence on tree production but an unclear influence on functions related to soil resource acquisition (Table 2, Figure 4), which partially confirms our first assumption on functional relationships (Figure 5a). In contrast to our conceptual model, there is limited evidence (from few studies) for a negative effect of fungi of a specific genetic identity and specific community assembly on root exudation (Table 2, 3).

The fungal properties in relation to soil exploration remain an understudied domain of the relationships between mycorrhizal fungal properties and tree function (Table 2). There is limited evidence for a positive effect of AMF spore density and AMF hyphal length on functions related to resource acquisition and tree production, but the influence of a specific ECMF exploration type is unclear. Each one study demonstrated a positive influence of hyphal foraging precision and a negative effect of the density of the ECM mycelium on tree production, which is the first confirmation of our second assumption on functional relationships (Figure 5b). The influence of fungal properties related to soil exploration on C release via root exudation or litter decomposition remains utterly unknown due to absence of investigation (Table 2, 3).

The influence of mycorrhizal colonization intensity on host C release from exudation and litter decomposition is unclear (Table 2), making it impossible to make a decision on our third assumption about functional relationships (Figure 5b). However, the positive influence of a specific mycorrhizal association type on root exudation and litter decomposition in combination with the negative influence of ECM tree dominance on litter decomposition may give an indirect hint of mycorrhizal control on tree functions related to C release (Table 2).

## 4 DISCUSSION

Mycorrhization is essential for basic tree functions, such as the release of root C and the acquisition of resources that promote the growth and fitness. According to the holobiont concept, the interaction between co-evolved host trees and root-associated endospheric fungi even has to be regarded a distinct ecological unit (Choi et al., 2021; Vandenkoornhuyse et al., 2015). While it is known that symbionts can co-construct host organs (Borges, 2017) and it has been hypothesized that genomic differences between mycorrhizal fungal guilds can reflect the eco-physiological traits of plant holobionts (Tedersoo & Bahram, 2019), functional analyses of holobionts – that is, analyses of mycorrhizal fungal properties in relation to host tree functions – have not yet been widely implemented. Recently, a comprehensive mycorrhizal trait framework has been developed that classifies traits based on their function and location among the symbiotic partners (Chaudhary et al., 2022). Here, we relate fungal properties and traits to host tree functions and summarize the demonstrated dependencies.

### 4.1 Important knowledge gaps in understanding the functions of holobionts of mycorrhizal trees

Based on our quantitative synthesis, we identify critical knowledge gaps to understand the functioning of mycorrhizal symbiosis in forest trees. These gaps originate from (i) temperate sampling bias and extremely limited knowledge from Afrotropical, dry-continental, and arctic tree or forest biomes, (ii) overrepresentation of five tree genera in trait analyses, of which three are phylogenetically closely related Fagaceae and only one genus (*Acer*) is widely used to study AM symbiosis of trees, (iii) underrepresentation of eco-physiological measurements of ERM and rare arbutoid mycorrhizas, and (iv) limited transfer of methods and studies to less controllable, natural field conditions with complex fungal communities and distant soil exploration by fungal hyphae. While the former three are probably the result of a Eurocentric and economic-targeted sampling bias, solving the latter is more complex since not all methods are applicable in the field.

Despite pleas from more than three decades ago (Read, 1991), the majority of current studies on mycorrhizal functions use inoculation with a limited number of easily culturable mycorrhizal fungal species to identify the influence of fungal genetic identity, diversity, or community composition on tree functioning (Bogar et al., 2022; Diagne et al., 2013; Thoen et al., 2020). Some studies stepped toward more natural conditions by growing host trees inoculated with a native pool of fungal species outdoors (Siqueira et al., 1998), or by growing trees with natural fungal assemblages under controlled experimental conditions (De Grandcourt et al., 2004; Köhler et al., 2018; Meier et al., 2013). Few studies have investigated the functions of forest trees associated with natural mycorrhizal fungal communities in forests (Anthony et al., 2022; Clemmensen et al., 2021; Khokon et al., 2023; Zavišić et al., 2018). Recently, mycorrhizal ecologists have been called to revisit old and new tools for studying the mechanistic functions of naturally complex fungal communities in the field (Lekberg & Helgason, 2018). These authors have identified several emerging new tools with potential for field application, including the production of manipulated mycorrhizal assemblages using *clustered regularly interspaced short palendromic repeats* (CRISPR) gene editing (Doudna & Charpentier, 2014); -omics approaches (e.g., transcriptomics, proteomics, and metabolomics) to study eco-physiological processes; and biomarker analyses (e.g., lab-on-a-chip microfluidics and quantum dots) to trace function. These new tools, when used alongside more established techniques such as mini-rhizotron imaging for hyphal dynamics and signature fatty acid analysis in combination with stable isotope labeling to determine plant carbon (C) allocation to mycelia, can be highly constructive (Lekberg & Helgason, 2018). For approaching the next level of mycorrhizal trait analysis in natural ecosystems, such exploration of new tools in combination with more established methods should be at the center of our interest. Taking advanced measurements of understudied mycorrhizal properties and traits will help identify those functional traits of holobionts that are critical for mycorrhizal symbiosis. These critical mycorrhizal traits should be included in trait databases on plant and fungal traits to improve our understanding of the role of mycorrhizal symbiosis in the functioning of global ecosystems.

### 4.2 ECMF exploration types can be vital for host tree functions

In our quantitative synthesis, we identify soil exploration as an understudied but potentially important mycorrhizal fungal property with great potential to understand tree functions. ECMFs can be ecologically classified by the distance from the root surface, which their emanating hyphae radiate into the soil, the presence and differentiation of their rhizomorphs, and the hydrophilicity of their hyphae (Agerer, 2001). These so-called exploration types represent different ECMF foraging strategies and are related to nutrient uptake, water transport rates, soil exploitation, soil organic matter formation and soil C cycling (Clemmensen et al., 2021; See et al., 2022). The morphology and chemistry of the extraradical mycelium can be used as vital traits of the ECMF, which are linked to functional variation between fungal taxa in the core of the mycorrhizal symbiosis (Agerer, 2001; Anthony et al., 2022; Lilleskov et al., 2011). In addition, different ECM exploration types have different abilities to adapt to N pollution in the forest ecosystem (Lilleskov et al., 2011; Suz et al., 2021), which could play a pivotal role in maintaining ecosystem functionality under the threat of global change. Surprisingly, few studies have directly related ECM exploration types to tree functioning so far. One study showed that the relative abundance of the medium-distance exploration type in the mycorrhizal fungal community affected tree N uptake (Pellitier et al., 2021). Another study demonstrated that the relative abundance of short- and medium-distance exploration types increased seedling survival (Jalón et al., 2020). Other authors found that the relative abundance of medium- and long-distance exploration types decreased tree growth, possibly as a consequence of greater C requirement for mycelium production (Anthony et al., 2022). As an alternative to exploration types, soil exploration can be represented by hyphal morphology (e.g., hyphal length and diameter), hyphal foraging precision and density, and the dispersal of rejuvenation organs (spore density). Several studies showed that these fungal propertys are positively related to the uptake and plant productivity (Chen et al., 2016; Sastry et al., 2000; Schoen et al., 2021). These promising results strengthen the potential importance of exploration type abundances for host tree functions, which requires further investigation.

### 4.3 Effects of mycorrhizal fungal community on host tree functions

Mycorrhizal fungal species can significantly differ in their effect on host tree function both intra- and interspecifically (Behm & Kiers, 2014; Johnson et al., 2012; van der Heijden et al., 2015) and in dependency on the fungal community assemblage (Pena & Polle, 2014). Differential functional trait responses were found between different fungal isolates of the same species (Hortal et al., 2017; Plett et al., 2015) and between individual fungal hyphae of a single species (Fernandez & Kennedy, 2018). The number of fungal species in a mycorrhizal community, i.e., local species co-existence, increases the chance of functional diversity, segregation among individual fungal species, and complementary mutualistic associations (Khokon et al., 2023). Our quantitative synthesis showed that the richness of more mycorrhizal fungal species enhances the acquisition of tree nutrients, water uptake, and plant productivity (Bompadre et al., 2013; Diagne et al., 2013; Köhler et al., 2018; Leberecht et al., 2016). Functional diversity in a mycorrhizal assemblage was positively related to the efficiency of P uptake efficiency in young beech trees (Köhler et al., 2018), while an opposite relationship was observed in mature beech ecosystems (Zavišić et al., 2018). The different effects of functional diversity are probably mediated by the inclusion of highly productive or pathogen species in the fungal community [selection effect (Meier et al., 2013)]. A complementary approach can be the measurement of functional genetic characteristics of mycorrhizal fungi and quantification of their expression level. This combination of transcriptomics with eco-physiological measurements can be powerful in identifying functional genetic variation between fungi in their effect on resource acquisition (Hortal et al., 2017), tree growth (Anthony et al., 2022), and decomposition processes (Pellitier & Zak, 2018). Likewise, the genetic evenness of fungal species in a community could be used as a measure of functional evenness, but this has rarely been carried out so far. However, it should be kept in mind that there are also functional differences between genes that are related to different trait categories. In a study in European forests, the proportion of genes that regulate energy and nutrient metabolism had a positive influence on tree growth, while organic N cycling genes had a negative effect (Anthony et al., 2022). Similarly, it remains unclear whether genes with saprotrophic function are actively transcribed when ECM taxa form mycorrhiza and whether their number can predict extracellular enzyme activity (Pellitier & Zak, 2018). Instead, they can also encode alternative plant-internal functions. Such molecular crosstalk between trees and mycorrhizal fungi is only beginning to be revealed (van der Heijden et al., 2015) and much progress can be made here.

### 4.4 Mycorrhizal control of root and litter C release

Although mycorrhizal colonization of tree roots can exert a strong control on root C release through exudation or decomposition, we are only just at the beginning of quantifying this effect and understanding the underlying processes. It was shown that the two major mycorrhizal types (i.e., ECM and AM) are functionally distinct in litter quality, enzymatic capacity, and C cost to host plants (Phillips et al., 2013; Smith & Read, 2010). This is reflected in, first, a greater loss of C with root exudation in ECM tree species (Liese et al., 2018; H. Yin et al., 2014) depending on root morphology (Akatsuki & Makita, 2020), and in, second, slower litter decomposition rates in ECM ecosystems (Carteron et al., 2022; Jiang et al., 2021; L. Yin et al., 2021), but faster decomposition by ECMF (Carteron et al., 2022). However, the question of whether the intensity of mycorrhizal colonization itself exerts control over root exudation and root litter decomposition remains contradictory (Argiroff et al., 2022; Meier et al., 2013; Trap et al., 2017). Furthermore, it has been debated whether a common mycorrhizal network between roots of multiple plants is used to shuttle C from one plant to another, but knowledge is currently still too scarce and unresolved to speculate on any control by the network on the root C release (Karst et al., 2023).

## 5 CONCLUSION

Our attempt to holistically examine mycorrhizal functions, setting fungal properties in relation to tree functions, has revealed several knowledge gaps arising from limited coverage of climate zones, tree species and mycorrhizal types. However, more critical than these sampling biases is the limited application of these methods in the field, where the most important mycorrhizal properties and traits for ecosystem functioning could be studied. We have yet to identify the fungal properties that are essential to the mycorrhizal symbiosis; these may not be the properties that are most frequently studied at present. Instead, these mycorrhizal properties are probably related to soil exploration, exploitation, and carbon (C) transfer. In order to assess functionally relevant mycorrhizal properties, a wide range of traits must be quantified in a diverse set of tree species and mycorrhizal types in different environments. Causal relationships with host functions must also be established. Our conceptual framework provides a mechanistic basis for identifying these fungal properties that have the strongest influence on host tree functions.

## ACKNOWLEDGEMENTS

The authors wish to thank Sandra Jacobi and Astrid Stilke of the Forestry and Timber Industry Library (Thünen Institute) for their excellent assistance in providing all non-digital publications. Ina C. Meier thanks the Deutsche Forschungsgemeinschaft (DFG, German Research Foundation) for financial support (Heisenberg program; grant no. ME 4156/5-1).

## AUTHOR CONTRIBUTIONS

ICM conceived the ideas and designed methodology; AMK collected the data; AMK analysed the data; AMK and ICM led the writing of the manuscript, contributed critically to the drafts, and gave final approval for publication.

## STATEMENT ON INCLUSION

Our study was a global review and was based on a meta-analysis of secondary data rather than primary data. As such, there was no local data collection.

## DATA AVAILABILITY STATEMENT

All data needed to evaluate the conclusions in the paper are present in the paper and/or the Supplementary Materials. All data analyzed in the study will be made publicly available as data publication after the participating researchers had the opportunity to publish the results. The R script including all analyses and figure preparations is available from the corresponding author upon reasonable request.

## DATA SOURCES

Our meta-analysis used data from multiple published resources which are provided in a separate Data Sources section.

## REFERENCES

Agerer, R. (2001). Exploration types of ectomycorrhizae. Mycorrhiza, 11(2), 107–114. 10.1007/s005720100108

Akatsuki, M., & Makita, N. (2020). Influence of fine root traits on in situ exudation rates in four conifers from different mycorrhizal associations. Tree Physiology, 40(8), 1071–1079. 10.1093/treephys/tpaa051

Anthony, M. A., Crowther, T. W., van der Linde, S., Suz, L. M., Bidartondo, M. I., Cox, F., Schaub, M., Rautio, P., Ferretti, M., Vesterdal, L., De Vos, B., Dettwiler, M., Eickenscheidt, N., Schmitz, A., Meesenburg, H., Andreae, H., Jacob, F., Dietrich, H.-P., Waldner, P., … Averill, C. (2022). Forest tree growth is linked to mycorrhizal fungal composition and function across Europe. The ISME Journal, 1–10. 10.1038/s41396-021-01159-7

Argiroff, W. A., Zak, D. R., Pellitier, P. T., Upchurch, R. A., & Belke, J. P. (2022). Decay by ectomycorrhizal fungi couples soil organic matter to nitrogen availability. Ecology Letters, 25(2), 391–404. 10.1111/ele.13923

Baxter, J. W., & Dighton, J. (2001). Ectomycorrhizal diversity alters growth and nutrient acquisition of grey birch (Betula populifolia) seedlings in host–symbiont culture conditions. New Phytologist, 152(1), 139–149. 10.1046/j.0028-646x.2001.00245.x

Behm, J. E., & Kiers, E. T. (2014). A phenotypic plasticity framework for assessing intraspecific variation in arbuscular mycorrhizal fungal traits. Journal of Ecology, 102(2), 315–327. 10.1111/1365-2745.12194

Bergmann, J., Weigelt, A., van der Plas, F., Laughlin, D. C., Kuyper, T. W., Guerrero-Ramirez, N., Valverde-Barrantes, O. J., Bruelheide, H., Freschet, G. T., Iversen, C. M., Kattge, J., McCormack, M. L., Meier, I. C., Rillig, M. C., Roumet, C., Semchenko, M., Sweeney, C. J., van Ruijven, J., York, L. M., & Mommer, L. (2020). The fungal collaboration gradient dominates the root economics space in plants. Science Advances, 6(27), eaba3756. 10.1126/sciadv.aba3756

Bogar, L. M., Tavasieff, O. S., Raab, T. K., & Peay, K. G. (2022). Does resource exchange in ectomycorrhizal symbiosis vary with competitive context and nitrogen addition? New Phytologist, 233(3), 1331–1344. 10.1111/nph.17871

Bompadre, M. J., Rios De Molina, M. C., Colombo, R. P., Fernandez Bidondo, L., Silvani, V. A., Pardo, A. G., Ocampo, J. A., & Godeas, A. M. (2013). Differential efficiency of two strains of the arbuscular mycorrhizal fungus Rhizophagus irregularis on olive (Olea europaea) plants under two water regimes. Symbiosis, 61(2), 105–112. 10.1007/s13199-013-0260-0

Borenstein, M., V. Hedges, L., P.T. Higgins, J., & R. Rothstein, H. (2009). Introduction to Meta-Analysis (1st ed.). John Wiley & Sons, Ltd. 10.1002/9780470743386

Borges, R. M. (2017). Co-niche construction between hosts and symbionts: Ideas and evidence. Journal of Genetics, 96(3), 483–489. 10.1007/s12041-017-0792-9

Brundrett, M. C. (2009). Mycorrhizal associations and other means of nutrition of vascular plants: Understanding the global diversity of host plants by resolving conflicting information and developing reliable means of diagnosis. Plant and Soil, 320(1), 37–77. 10.1007/s11104-008-9877-9

Brundrett, M., & Tedersoo, L. (2019). Misdiagnosis of mycorrhizas and inappropriate recycling of data can lead to false conclusions. New Phytologist, 221(1), 18–24. 10.1111/nph.15440

Carteron, A., Cichonski, F., & Laliberté, E. (2022). Ectomycorrhizal Stands Accelerate Decomposition to a Greater Extent than Arbuscular Mycorrhizal Stands in a Northern Deciduous Forest. Ecosystems, 25(6), 1234–1248. 10.1007/s10021-021-00712-x

Chaudhary, V. B., Holland, E. P., Charman-Anderson, S., Guzman, A., Bell-Dereske, L., Cheeke, T. E., Corrales, A., Duchicela, J., Egan, C., Gupta, M. M., Hannula, S. E., Hestrin, R., Hoosein, S., Kumar, A., Mhretu, G., Neuenkamp, L., Soti, P., Xie, Y., & Helgason, T. (2022). What are mycorrhizal traits? Trends in Ecology & Evolution. 10.1016/j.tree.2022.04.003

Chaudhary, V. B., Rúa, M. A., Antoninka, A., Bever, J. D., Cannon, J., Craig, A., Duchicela, J., Frame, A., Gardes, M., Gehring, C., Ha, M., Hart, M., Hopkins, J., Ji, B., Johnson, N. C., Kaonongbua, W., Karst, J., Koide, R. T., Lamit, L. J., … Hoeksema, J. D. (2016). MycoDB, a global database of plant response to mycorrhizal fungi. Scientific Data, 3(1), Article 1. 10.1038/sdata.2016.28

Chen, W., Koide, R. T., Adams, T. S., DeForest, J. L., Cheng, L., & Eissenstat, D. M. (2016). Root morphology and mycorrhizal symbioses together shape nutrient foraging strategies of temperate trees. Proceedings of the National Academy of Sciences, 113(31), 8741–8746. 10.1073/pnas.1601006113

Choi, K., Khan, R., & Lee, S.-W. (2021). Dissection of plant microbiota and plant-microbiome interactions. Journal of Microbiology, 59(3), 281–291. 10.1007/s12275-021-0619-5

Clemmensen, K. E., Bahr, A., Ovaskainen, O., Dahlberg, A., Ekblad, A., Wallander, H., Stenlid, J., Finlay, R. D., Wardle, D. A., & Lindahl, B. D. (2013). Roots and Associated Fungi Drive Long-Term Carbon Sequestration in Boreal Forest. Science, 339(6127), 1615–1618. 10.1126/science.1231923

Clemmensen, K. E., Durling, M. B., Michelsen, A., Hallin, S., Finlay, R. D., & Lindahl, B. D. (2021). A tipping point in carbon storage when forest expands into tundra is related to mycorrhizal recycling of nitrogen. Ecology Letters, 24(6), 1193–1204. 10.1111/ele.13735

Cornelissen, J., Aerts, R., Cerabolini, B., Werger, M., & van der Heijden, M. (2001). Carbon cycling traits of plant species are linked with mycorrhizal strategy. Oecologia, 129(4), 611–619. 10.1007/s004420100752

Dawson, S. K., Carmona, C. P., González-Suárez, M., Jönsson, M., Chichorro, F., Mallen-Cooper, M., Melero, Y., Moor, H., Simaika, J. P., & Duthie, A. B. (2021). The traits of “trait ecologists”: An analysis of the use of trait and functional trait terminology. Ecology and Evolution, 11(23), 16434–16445. 10.1002/ece3.8321

de Bello, F., Lavorel, S., Díaz, S., Harrington, R., Cornelissen, J. H. C., Bardgett, R. D., Berg, M. P., Cipriotti, P., Feld, C. K., Hering, D., Martins da Silva, P., Potts, S. G., Sandin, L., Sousa, J. P., Storkey, J., Wardle, D. A., & Harrison, P. A. (2010). Towards an assessment of multiple ecosystem processes and services via functional traits. Biodiversity and Conservation, 19(10), 2873–2893. 10.1007/s10531-010-9850-9

De Grandcourt, A., Epron, D., Montpied, P., Louisanna, E., Béreau, M., Garbaye, J., & Guehl, J.-M. (2004). Contrasting responses to mycorrhizal inoculation and phosphorus availability in seedlings of two tropical rainforest tree species. New Phytologist, 161(3), 865–875. 10.1046/j.1469-8137.2004.00978.x

Diagne, N., Thioulouse, J., Sanguin, H., Prin, Y., Krasova-Wade, T., Sylla, S., Galiana, A., Baudoin, E., Neyra, M., Svistoonoff, S., Lebrun, M., & Duponnois, R. (2013). Ectomycorrhizal diversity enhances growth and nitrogen fixation of *Acacia mangium* seedlings. Soil Biology and Biochemistry, 57, 468–476. 10.1016/j.soilbio.2012.08.030

Doudna, J. A., Charpentier, E. (2014) The new frontier of genome engineering with CRISPR-Cas9. Science, 346, 6213. 10.1126/science.l25809

Egger, M., Davey Smith, G., Schneider, M., & Minder, C. (1997). Bias in meta-analysis detected by a simple, graphical test. BMJ (Clinical Research Ed.), 315(7109), 629–634. 10.1136/bmj.315.7109.629

Fanin, N., Clemmensen, K. E., Lindahl, B. D., Farrell, M., Nilsson, M.-C., Gundale, M. J., Kardol, P., & Wardle, D. A. (2022). Ericoid shrubs shape fungal communities and suppress organic matter decomposition in boreal forests. New Phytologist, 236(2), 684–697. 10.1111/nph.18353

Fernandez, C. W., & Kennedy, P. G. (2018). Melanization of mycorrhizal fungal necromass structures microbial decomposer communities. Journal of Ecology, 106(2), 468–479. 10.1111/1365-2745.12920

Garnier, S., Ross, N., Rudis, boB, Filipovic-Pierucci, A., Galili, T., timelyportfolio, Greenwell, B., Sievert, C., Harris, D. J., Sciaini, M., & Chen, J. J. (2021). sjmgarnier/viridis: CRAN release v0.6.2 [Computer software]. Zenodo. 10.5281/zenodo.5579397

Halbritter, A. H., De Boeck, H. J., Eycott, A. E., Reinsch, S., Robinson, D. A., Vicca, S., Berauer, B., Christiansen, C. T., Estiarte, M., Grünzweig, J. M., Gya, R., Hansen, K., Jentsch, A., Lee, H., Linder, S., Marshall, J., Peñuelas, J., Kappel Schmidt, I., Stuart-Haëntjens, E., … Vandvik, V. (2020). The handbook for standardized field and laboratory measurements in terrestrial climate change experiments and observational studies (ClimEx). Methods in Ecology and Evolution, 11(1), 22–37. 10.1111/2041-210X.13331

Hazard, C., & Johnson, D. (2018). Does genotypic and species diversity of mycorrhizal plants and fungi affect ecosystem function? New Phytologist, 220(4), 1122–1128. 10.1111/nph.15010

Hedges, L. V., & Olkin, I. (1985). Statistical Methods for Meta-Analysis. https://shop.elsevier.com/books/statistical-methods-for-meta-analysis/hedges/978-0-08-057065-5

Hodge, A., & Fitter, A. H. (2010). Substantial nitrogen acquisition by arbuscular mycorrhizal fungi from organic material has implications for N cycling. Proceedings of the National Academy of Sciences, 107(31), 13754–13759. 10.1073/pnas.1005874107

Hortal, S., Plett, K. L., Plett, J. M., Cresswell, T., Johansen, M., Pendall, E., & Anderson, I. C. (2017). Role of plant–fungal nutrient trading and host control in determining the competitive success of ectomycorrhizal fungi. The ISME Journal, 11(12), 2666–2676. 10.1038/ismej.2017.116

Iversen, C. M., McCormack, M. L., Powell, A. S., Blackwood, C. B., Freschet, G. T., Kattge, J., Roumet, C., Stover, D. B., Soudzilovskaia, N. A., Valverde-Barrantes, O. J., van Bodegom, P. M., & Violle, C. (2017). A global Fine-Root Ecology Database to address below-ground challenges in plant ecology. New Phytologist, 215(1), 15–26. 10.1111/nph.14486

Jalón, L. G. de, Limousin, J.-M., Richard, F., Gessler, A., Peter, M., Hättenschwiler, S., & Milcu, A. (2020). Microhabitat and ectomycorrhizal effects on the establishment, growth and survival of Quercus ilex L. seedlings under drought. PLOS ONE, 15(6), e0229807. 10.1371/journal.pone.0229807

Jiang, L., Wang, H., Li, S., Fu, X., Dai, X., Yan, H., & Kou, L. (2021). Mycorrhizal and environmental controls over root trait–decomposition linkage of woody trees. New Phytologist, 229(1), 284–295. 10.1111/nph.16844

Jin, Y., & Qian, H. (2019). V.PhyloMaker: An R package that can generate very large phylogenies for vascular plants. Ecography, 42(8), 1353–1359. 10.1111/ecog.04434

Johnson, D., Martin, F., Cairney, J. W. G., & Anderson, I. C. (2012). The importance of individuals: Intraspecific diversity of mycorrhizal plants and fungi in ecosystems. New Phytologist, 194(3), 614–628. 10.1111/j.1469-8137.2012.04087.x

Karst, J., Jones, M. D., & Hoeksema, J. D. (2023). Positive citation bias and overinterpreted results lead to misinformation on common mycorrhizal networks in forests. Nature Ecology & Evolution, 1–11. 10.1038/s41559-023-01986-1

Khokon, A. M., Janz, D., & Polle, A. (2023). Ectomycorrhizal diversity, taxon-specific traits and root N uptake in temperate beech forests. New Phytologist, 239(2), 739–751. 10.1111/nph.18978

Köhler, J., Yang, N., Pena, R., Raghavan, V., Polle, A., & Meier, I. C. (2018). Ectomycorrhizal fungal diversity increases phosphorus uptake efficiency of European beech. New Phytologist, 220(4), 1200–1210. 10.1111/nph.15208

Leberecht, M., Tu, J., & Polle, A. (2016). Acid and calcareous soils affect nitrogen nutrition and organic nitrogen uptake by beech seedlings (Fagus sylvatica L.) under drought, and their ectomycorrhizal community structure. Plant and Soil, 409(1), 143–157. 10.1007/s11104-016-2956-4

Lekberg, Y., & Helgason, T. (2018). In situ mycorrhizal function – knowledge gaps and future directions. New Phytologist, 220(4), 957–962. 10.1111/nph.15064

Leuschner, C., & Meier, I. C. (2018). The ecology of Central European tree species: Trait spectra, functional trade-offs, and ecological classification of adult trees. Perspectives in Plant Ecology, Evolution and Systematics, 33, 89–103. 10.1016/j.ppees.2018.05.003

Liese, R., Alings, K., & Meier, I. C. (2017). Root Branching Is a Leading Root Trait of the Plant Economics Spectrum in Temperate Trees. Frontiers in Plant Science, 8. https://www.frontiersin.org/articles/10.3389/fpls.2017.00315

Liese, R., Lübbe, T., Albers, N. W., & Meier, I. C. (2018). The mycorrhizal type governs root exudation and nitrogen uptake of temperate tree species. Tree Physiology, 38(1), 83–95. 10.1093/treephys/tpx131

Lilleskov, E. A., Hobbie, E. A., & Horton, T. R. (2011). Conservation of ectomycorrhizal fungi: Exploring the linkages between functional and taxonomic responses to anthropogenic N deposition. Fungal Ecology, 4(2), 174–183. 10.1016/j.funeco.2010.09.008

Lilleskov, E. A., Kuyper, T. W., Bidartondo, M. I., & Hobbie, E. A. (2019). Atmospheric nitrogen deposition impacts on the structure and function of forest mycorrhizal communities: A review. Environmental Pollution, 246, 148–162. 10.1016/j.envpol.2018.11.074

Marro, N., Grilli, G., Soteras, F., Caccia, M., Longo, S., Cofré, N., Borda, V., Burni, M., Janoušková, M., Urcelay, C. (2022) The effects of arbuscular mycorrhizal fungal species and taxonomic groups on stressed and unstressed plants: a global meta-analysis. New Phytologist, 235, 320–332. 10.1111/nph.18102

McCormack, M. L., Iversen, C. M. (2019) Physical and functional constraints on viable belowground acquisition strategies. Frontiers in Plant Science, 10, 1215. 10.3389/fpls.2019.01215

Meier, I. C., Avis, P. G., & Phillips, R. P. (2013). Fungal communities influence root exudation rates in pine seedlings. FEMS Microbiology Ecology, 83(3), 585–595. 10.1111/1574-6941.12016

Moher, D., Liberati, A., Tetzlaff, J., & Altman, D. G. (2010). Preferred reporting items for systematic reviews and meta-analyses: The PRISMA statement. International Journal of Surgery, 8(5), 336–341. 10.1016/j.ijsu.2010.02.007

Nguyen, N. H., Song, Z., Bates, S. T., Branco, S., Tedersoo, L., Menke, J., Schilling, J. S., & Kennedy, P. G. (2016). FUNGuild: An open annotation tool for parsing fungal community datasets by ecological guild. Fungal Ecology, 20, 241–248. 10.1016/j.funeco.2015.06.006

Page, M. J., McKenzie, J. E., Bossuyt, P. M., Boutron, I., Hoffmann, T. C., Mulrow, C. D., Shamseer, L., Tetzlaff, J. M., Akl, E. A., Brennan, S. E., Chou, R., Glanville, J., Grimshaw, J. M., Hróbjartsson, A., Lalu, M. M., Li, T., Loder, E. W., Mayo-Wilson, E., McDonald, S., McGuiness, L. A., Stewart, L. A., Thomas, J., Tricco, A. C., Welch, V. A., Whiting, P., Moher, D. (2021) The PRISMA 2020 statement: an updated guideline for reporting systematic reviews. BMJ, 372, 71. 10.1136/bmj.n71

Pellitier, P. T., & Zak, D. R. (2018). Ectomycorrhizal fungi and the enzymatic liberation of nitrogen from soil organic matter: Why evolutionary history matters. New Phytologist, 217(1), 68–73. 10.1111/nph.14598

Pellitier, P. T., Zak, D. R., Argiroff, W. A., & Upchurch, R. A. (2021). Coupled Shifts in Ectomycorrhizal Communities and Plant Uptake of Organic Nitrogen Along a Soil Gradient: An Isotopic Perspective. Ecosystems, 24(8), 1976–1990. 10.1007/s10021-021-00628-6

Pena, R., & Polle, A. (2014). Attributing functions to ectomycorrhizal fungal identities in assemblages for nitrogen acquisition under stress. The ISME Journal, 8(2), Article 2. 10.1038/ismej.2013.158

Phillips, R. P., Brzostek, E., & Midgley, M. G. (2013). The mycorrhizal-associated nutrient economy: A new framework for predicting carbon–nutrient couplings in temperate forests. New Phytologist, 199(1), 41–51. 10.1111/nph.12221

Plett, J. M., Kohler, A., Khachane, A., Keniry, K., Plett, K. L., Martin, F., & Anderson, I. C. (2015). The effect of elevated carbon dioxide on the interaction between Eucalyptus grandis and diverse isolates of Pisolithus sp. Is associated with a complex shift in the root transcriptome. New Phytologist, 206(4), 1423–1436. 10.1111/nph.13103

Põlme, S., Abarenkov, K., Henrik Nilsson, R., Lindahl, B. D., Clemmensen, K. E., Kauserud, H., Nguyen, N., Kjøller, R., Bates, S. T., Baldrian, P., Frøslev, T. G., Adojaan, K., Vizzini, A., Suija, A., Pfister, D., Baral, H.-O., Järv, H., Madrid, H., Nordén, J., … Tedersoo, L. (2020). FungalTraits: A user-friendly traits database of fungi and fungus-like stramenopiles. Fungal Diversity, 105(1), 1–16. 10.1007/s13225-020-00466-2

R development Core Team. (2020). R: The R Project for Statistical Computing: R Foundation for Statistical Computing: Vienna, Austria, 2020. https://www.r-project.org/

Read, D. J. (1991). Mycorrhizas in ecosystems. Experientia, 47(4), 376–391. 10.1007/BF01972080

Read, D. J., & Perez-Moreno, J. (2003). Mycorrhizas and nutrient cycling in ecosystems – a journey towards relevance? New Phytologist, 157(3), 475–492. 10.1046/j.1469-8137.2003.00704.x

Rosenberg, M. S. (2005). The File-Drawer Problem Revisited: A General Weighted Method for Calculating Fail-Safe Numbers in Meta-Analysis. Evolution, 59(2), 464–468. 10.1111/j.0014-3820.2005.tb01004.x

Rosenthal, R. (1991). Meta-analysis: A review. Biopsychosocial Science and Medicine, 53(3), 247.

Sastry, M. S. R., Sharma, A. K., & Johri, B. N. (2000). Effect of an AM fungal consortium and Pseudomonas on the growth and nutrient uptake of Eucalyptus hybrid. Mycorrhiza, 10(2), 55–61. 10.1007/s005720000057

Schoen, C., Montibeler, M., Costa, M. D., Antunes, P. M., & Stürmer, S. L. (2021). Inter and intra-specific variability in arbuscular mycorrhizal fungi affects hosts and soil health. Symbiosis, 85(3), 273–289. 10.1007/s13199-021-00812-1

See, C. R., Keller, A. B., Hobbie, S. E., Kennedy, P. G., Weber, P. K., & Pett-Ridge, J. (2022). Hyphae move matter and microbes to mineral microsites: Integrating the hyphosphere into conceptual models of soil organic matter stabilization. Global Change Biology, 28(8), 2527–2540. 10.1111/gcb.16073

Siqueira, J. O., Saggin-Júnior, O. J., Flores-Aylas, W. W., & Guimarães, P. T. G. (1998). Arbuscular mycorrhizal inoculation and superphosphate application influence plant development and yield of coffee in Brazil. Mycorrhiza, 7(6), 293–300. 10.1007/s005720050195

Smith, S. E., & Read, D. J. (2010). Mycorrhizal Symbiosis. Academic Press.

Soudzilovskaia, N. A., Douma, J. C., Akhmetzhanova, A. A., van Bodegom, P. M., Cornwell, W. K., Moens, E. J., Treseder, K. K., Tibbett, M., Wang, Y.-P., & Cornelissen, J. H. C. (2015). Global patterns of plant root colonization intensity by mycorrhizal fungi explained by climate and soil chemistry. Global Ecology and Biogeography, 24(3), 371–382. 10.1111/geb.12272

Soudzilovskaia, N. A., Vaessen, S., Barcelo, M., He, J., Rahimlou, S., Abarenkov, K., Brundrett, M. C., Gomes, S. I. F., Merckx, V., & Tedersoo, L. (2020). FungalRoot: Global online database of plant mycorrhizal associations. New Phytologist, 227(3), 955–966. 10.1111/nph.16569

Steidinger, B. S., Crowther, T. W., Liang, J., Van Nuland, M. E., Werner, G. D. A., Reich, P. B., Nabuurs, G. J., de-Miguel, S., Zhou, M., Picard, N., Herault, B., Zhao, X., Zhang, C., Routh, D., & Peay, K. G. (2019). Climatic controls of decomposition drive the global biogeography of forest-tree symbioses. Nature, 569(7756), Article 7756. 10.1038/s41586-019-1128-0

Suz, L. M., Bidartondo, M. I., van der Linde, S., & Kuyper, T. W. (2021). Ectomycorrhizas and tipping points in forest ecosystems. New Phytologist, 231(5), 1700–1707. 10.1111/nph.17547

Tang, B., Man, J., Lehmann, A., Rillig, M. C. (2023) Arbuscular mycorrhizal fungi benefit plants in response to major global change factors. Ecology Letters, 26, 2087–2097. 10.1111/ele.14320

Tedersoo, L., & Bahram, M. (2019). Mycorrhizal types differ in ecophysiology and alter plant nutrition and soil processes. Biological Reviews, 94(5), 1857–1880. 10.1111/brv.12538

Thoen, E., Harder, C. B., Kauserud, H., Botnen, S. S., Vik, U., Taylor, A. F. S., Menkis, A., & Skrede, I. (2020). In vitro evidence of root colonization suggests ecological versatility in the genus Mycena. New Phytologist, 227(2), 601–612. 10.1111/nph.16545

Trap, J., Akpa-Vinceslas, M., Margerie, P., Boudsocq, S., Richard, F., Decaëns, T., & Aubert, M. (2017). Slow decomposition of leaf litter from mature Fagus sylvatica trees promotes offspring nitrogen acquisition by interacting with ectomycorrhizal fungi. Journal of Ecology, 105(2), 528–539. 10.1111/1365-2745.12665

Van der Heijden, M. G. A., Martin, F. M., Selosse, M.-A., & Sanders, I. R. (2015). Mycorrhizal ecology and evolution: The past, the present, and the future. New Phytologist, 205(4), 1406–1423. 10.1111/nph.13288

Vandenkoornhuyse, P., Quaiser, A., Duhamel, M., Van, A. L., & Dufresne, A. (2015). The importance of the microbiome of the plant holobiont. New Phytologist, 206(4), 1196–1206. 10.1111/nph.13312

Větrovský, T., Morais, D., Kohout, P., Lepinay, C., Algora, C., Awokunle Hollá, S., Bahnmann, B. D., Bílohnědá, K., Brabcová, V., D’Alò, F., Human, Z. R., Jomura, M., Kolařík, M., Kvasničková, J., Lladó, S., López-Mondéjar, R., Martinović, T., Mašínová, T., Meszárošová, L., … Baldrian, P. (2020). GlobalFungi, a global database of fungal occurrences from high-throughput-sequencing metabarcoding studies. Scientific Data, 7(1), Article 1. 10.1038/s41597-020-0567-7

Viechtbauer, W. (2010). Conducting Meta-Analyses in R with the metafor Package. Journal of Statistical Software, 36, 1–48. 10.18637/jss.v036.i03

Violle, C., Navas, M.-L., Vile, D., Kazakou, E., Fortunel, C., Hummel, I., & Garnier, E. (2007). Let the concept of trait be functional! Oikos, 116(5), 882–892. 10.1111/j.0030-1299.2007.15559.x

Wan, X., Wang, W., Liu, J., & Tong, T. (2014). Estimating the sample mean and standard deviation from the sample size, median, range and/or interquartile range. BMC Medical Research Methodology, 14(1), 135. 10.1186/1471-2288-14-135

Wickham, H. (2009). ggplot2: Elegant Graphics for Data Analysis. Springer-Verlag. 10.1007/978-0-387-98141-3

Wickham, H., Averick, M., Bryan, J., Chang, W., McGowan, L. D., François, R., Grolemund, G., Hayes, A., Henry, L., Hester, J., Kuhn, M., Pedersen, T. L., Miller, E., Bache, S. M., Müller, K., Ooms, J., Robinson, D., Seidel, D. P., Spinu, V., … Yutani, H. (2019). Welcome to the Tidyverse. Journal of Open Source Software, 4(43), 1686. 10.21105/joss.01686

Wright, I. J., Reich, P. B., Westoby, M., Ackerly, D. D., Baruch, Z., Bongers, F., Cavender-Bares, J., Chapin, T., Cornelissen, J. H. C., Diemer, M., Flexas, J., Garnier, E., Groom, P. K., Gulias, J., Hikosaka, K., Lamont, B. B., Lee, T., Lee, W., Lusk, C., … Villar, R. (2004). The worldwide leaf economics spectrum. Nature, 428(6985), Article 6985. 10.1038/nature02403

Yin, H., Wheeler, E., & Phillips, R. P. (2014). Root-induced changes in nutrient cycling in forests depend on exudation rates. Soil Biology and Biochemistry, 78, 213–221. 10.1016/j.soilbio.2014.07.022

Yin, L., Dijkstra, F. A., Phillips, R. P., Zhu, B., Wang, P., & Cheng, W. (2021). Arbuscular mycorrhizal trees cause a higher carbon to nitrogen ratio of soil organic matter decomposition via rhizosphere priming than ectomycorrhizal trees. Soil Biology and Biochemistry, 157, 108246. 10.1016/j.soilbio.2021.108246

Yu, G., Smith, D. K., Zhu, H., Guan, Y., & Lam, T. T.-Y. (2017). ggtree: An r package for visualization and annotation of phylogenetic trees with their covariates and other associated data. Methods in Ecology and Evolution, 8(1), 28–36. 10.1111/2041-210X.12628

Zanne, A. E., Abarenkov, K., Afkhami, M. E., Aguilar-Trigueros, C. A., Bates, S., Bhatnagar, J. M., Busby, P. E., Christian, N., Cornwell, W., Crowther, T. W., Moreno, H. F., Floudas, D., Gazis, R., Hibbett, D., Kennedy, P., Lindner, D. L., Maynard, D. S., Milo, A. M., Nilsson, H., … Treseder, K. K. (2019). Fungal functional ecology: Bringing a trait-based approach to plant-associated fungi. EcoEvoRxiv. 10.32942/osf.io/a7f6g

Zavišić, A., Yang, N., Marhan, S., Kandeler, E., & Polle, A. (2018). Forest Soil Phosphorus Resources and Fertilization Affect Ectomycorrhizal Community Composition, Beech P Uptake Efficiency, and Photosynthesis. Frontiers in Plant Science, 9. https://www.frontiersin.org/articles/10.3389/fpls.2018.00463

